# Brain transcriptomics reveals features of the host response after dengue virus type I infection induced central nervous system injury in AGB6 mice

**DOI:** 10.1101/2024.01.10.574956

**Authors:** Ning Yu, Shigang Chen, Yumeng Liu, Peng Wang, Longlong Wang, Ningning Hu, He Zhang, Xiao Li, Huijun Lu, Ningyi Jin

## Abstract

Dengue viruses are the causative agents of dengue fever, dengue hemorrhagic fever, and dengue shock syndrome, which are mainly transmitted by Aedes aegypti and Aedes albopictus mosquitoes, and cost billions of dollars annually in medical and mosquito control. It is estimated that up to 20% of dengue virus infections affect the brain. The incidence of DENV infection of the nervous system is increasing, suggesting that more people are at risk for neurological complications of the infection. Currently, understanding the pathogenic mechanisms of dengue virus infection of the nervous system is hampered by the lack of suitable small animal models. Here, we analyzed the pathogenicity of dengue virus in type I and type Ⅱ interferon receptor-deficient mice by intraperitoneal inoculation of DENV-I. Infected mice showed such neurological symptoms as opisthotonus, hunching, ataxia, and paralysis of one or both hind limbs. Viremia can be detected 3 days after infection. The virus showed evident brain tropism post-inoculation and viral loads peaked at 14 days post-inoculation in infected mice brain. Significant histopathologic changes were observed in brain tissue (hippocampal region and cerebral cortex). Immunohistochemistry and fluorescence showed clear fluorescent signals. Hematological analysis showed hemorrhage and hemoconcentration in DENV-I infected mice. Subsequently, brain tissue transcriptome sequencing was performed to assess host response characteristics in DENV-I infected AGB6 mice. Functional enrichment analysis and analysis of significantly differentially expressed genes (DEGs) were used to identify the key molecular mechanisms of brain tissue injury. Transcriptional patterns in brain tissue suggest that aberrant expression of pro-inflammatory cytokines induces antiviral responses to dengue virus and tissue damage. Combined with the KEGG results, it was hypothesized that the probable cause of the neurological damage affecting the neurological system and hindlimb paralysis was the marked enrichment of upregulated DEGs in the cytokine-cytokine receptor interaction signaling pathway. Screening of hub genes and their characterization by qPCR and ELISA, it was hypothesized that IL-6 and IFN-γ might be the key factors in dengue virus-induced neurological manifestations.

## Introduction

Dengue virus belongs to the *Flaviviridae* family, *Flavivirus* genus, a single-stranded positive-stranded RNA virus, which is a mosquito-borne virus like West Nile virus, Zika virus, and yellow fever virus[1]. The genome of this virus has only one open reading frame (ORF), which is translated into a multimeric protein, which is subsequently broken down into three structural and seven nonstructural proteins catalyzed by host proteases and the virus’s protease, and performs their respective functions. The order of the encoded proteins is 5’-C-PrM-E-NS1-NS2a-NS2b-NS3-NS4a-NS4b-NS5-3’[2, 3]. According to their surface antigenicity, dengue viruses are classified into four serotypes DENV-I, DENV-II, DENV-III, and DENV-IV[4]. DENV-I was first detected in 1943 in French Polynesia and Japan and seemed to be the most reported serotype between 1943 and 2013[5, 6]. Dengue fever is a self-limiting disease; most people with dengue have no or mild symptoms, and severe cases can cause dengue hemorrhagic fever (DHF) and dengue shock syndrome (DSS) [7, 8]. Dengue fever is expected to infect an estimated 100 million to 400 million people each year[9], posing a significant threat to the global economy and health.

Dengue viruses were earlier misidentified as non-neurophilic, and in 2009, the WHO released new guidelines and classifications for dengue, which included CNS involvement in the definition of severe disease[10]. Manifestations of meningitis, encephalitis, Guillain-Barre syndrome (GBS), acute demyelinating encephalopathy (ADEM), myelitis, polyneuropathy, and mononeuropathy[11]. The incidence of encephalopathy and encephalitis, the most common neurological complications of dengue, has been estimated to be between 0.5 and 6.2%[12]. Currently, the pathogenesis of dengue neurologic disorders is incompletely understood. Three mechanisms may be operative: direct CNS invasion by the virus, autoimmune reactions, and metabolic alterations[13].

Transcriptome sequencing enables the study of gene function and gene structure at a holistic level, revealing the molecular mechanisms involved in specific biological processes as well as in the development of diseases. Clinical data have demonstrated a significant relationship between brain tissue damage and severe dengue fever[14–16]. Currently, there is no breakthrough in the study of the pathogenic mechanism of dengue virus-induced encephalitis, and there is still a need to explore the general features of host inflammatory and immune responses in DENV-infected mouse models, especially the transcriptional responses in brain injury, which may reveal potential therapeutic targets from a holistic perspective.

In this study, we aimed to assess the transcriptomic profile of brain tissue in a mouse model of dengue virus infection based on a lethal animal model, followed by a series of functional analyses to partially elucidate the underlying pathogenesis of dengue virus-induced brain damage.

## Materials and methods

### Cell culture

The African green monkey kidney cells (Vero) were cultured in DMEM medium supplemented with 10% FBS. An *Aedes albopictus* gut line (C6/36) was purchased from the National Collection of Authenticated Cell Cultures. C6/36 cells were cultured in MEM medium with 15% fetal bovine serum (FBS) and incubated in a humidified atmosphere containing 5% CO_2_ at 28℃.

### Virus strain

The virus strain used in the study derives from a patient in RuiLi, YunNan Province in 2017 that has been exclusively passaged for about 6 rounds in C6/36 cells (GenBank number: OM250394). For AGB6 infection, viruses were harvested and concentrated using a 10K MWCO Amicon (Merck) at 1500×g, 4℃, for 2 hours, then quantified by immunoperoxidase monolayer assays in Vero cells. Virus titer after concentration was 10^5.63^/0.1 mL.

### Mice

Male AGB6 mice (deficient in both type I and type II interferon receptor) were a cross between *Ifnar^-/-^* mice (C57BL/6 background) and *Ifngr^-/-^*mice (C57BL/6 background) were purchased from Shanghai Model Organisms Center, Inc. Female C57BL/6 and BALB/c were purchased from Beijing Viton Lihua Laboratory Animal Technology Co. They were housed in the Animal Biological Safety Level-2 laboratory (ABSL-2) at Changchun Veterinary Research Institute and were provided access to food and water. Unless otherwise specified, both male and female mice between the ages of 5-6 weeks were used in the study.

### Mice infection

To test the virulence of the DENV-I, twenty-five AGB6 mice were randomly divided to five groups. They were administered with 10^8^ to 10^5^ TCID_50_ of DENV-I via the intraperitoneal (IP.) route (0.5 mL in sterile PBS). Mice injected with an equivalent volume of PBS (mock infection) were used as the negative control. Animals were monitored for clinical phenotypes, morbidity, and mortality. For the 10^7^ TCID_50_ infected group. At 3, 7, and 11 dpi (n=3), blood was taken from the tail vein to determine viremia, then mice were euthanized with tribromoethanol, and the tissues (heart, liver, spleen, lung, kidney, brain, jejunum, ileum, caecum, colon, rectum) were quickly homogenized on ice and stored at -40℃ for further use. When a humane endpoint (body weight loss of ≥20%, hunched posture, ruffled fur, conjunctivitis, movement impairment, lower limb paralysis) was reached, mice (n=3) were euthanized, the half of brain was collected and quick-frozen in liquid nitrogen, and then RNA was extracted for transcriptome sequencing.

To clarify the infection of DENV-I in immunocompetent mice. C57BL/6 (n=12) and BALB/c (n=12) mice were administered with 10^7^ TCID_50_ of DENV-I via the intraperitoneal (IP.) route (0.5 mL of sterile PBS). They were weighed and the course of the disease was monitored until 15 dpi. Blood and tissues (n=3) were collected at the same intervals selected for the infected group of AGB6 mice (10^7^ TCID_50_) and stored at -40 for subsequent analysis.

### Quantitative real-time (qRT) PCR

The total RNA was extracted from the blood, heart, liver, spleen, lung, kidney, brain, jejunum, ileum, caecum, colon, and rectum (obtained from AGB6 mice, C57BL/6 mice and BALB/c mice) using a Total RNA Extractor kit (Sangon Biotech, China). Using the HiScript II U+ One Step qRT-PCR Probe Kit (Vazyme, Nanjing), 150 μg of RNA was pipetted for amplification. Using previously established assays, serial dilutions of the pUC57-DENV plasmid with known concentrations were used to establish the standard curve. RNA copies per mL or RNA copies per gram of each sample were calculated from the Ct values.

The M-MLV Reverse Transcriptase Kit was used to reverse-transcribe the total RNA to cDNA. The relative expression levels of IL-1β, IL-6, IFN-γ, TNFα, CCL2, CCL3, CCL4, and CCL5 were determined using RT-PCR. Mouse GAPDH internal reference primers were purchased from Sangon Biotech (Shanghai) Co., Ltd. The primers used in this study are listed in Supplementary Table S1.

### Viral protein detection

For Western blot analysis, 10 μL of tissue slurry samples were run on a 10%-15% Bis-Tris SDS-PAGE gel as described previously[17]. The Nitrocellulose (NC) membranes were incubated with Anti-DENV-E gene (Abcam, ab9202) and NS1 gene (Genetex, GTX124280) antibodies. Horseradish peroxidase (HRP) secondary antibody (Beyotime, A0216) was used at a 1:2000 dilution. NC membranes were imaged on an Amersham Imager 600 (General Electric, USA).

### Antibody titers

Systemic antibody titers against DENV-I were determined by Enzyme-linked Immunosorbent Assays (ELISA) as described previously[18]. Briefly, 96-well plates were coated overnight at 4℃ with 10^5^ TCID_50_/0.1mL of heat-inactivated DENV-I in 0.1M NaHCO_3_ buffer at pH 9.6. Two-fold serially diluted serum samples (1:50 to 1:102,400) were added to the wells and incubated for 2 hours at 37℃. HRP-conjugated anti-mouse IgM (Abcam, ab97230) and IgG (H+L) (Abcam, ab6808) secondary antibodies were used. The ELISA titers were defined as the reciprocal of the highest serum dilution that equals to 3 times the absorbance reading from uninfected mouse serum sample.

### Histopathology Staining

Paraffin-embedded tissue sections (slice thickness of 4μm) were dewaxed with xylol, hydrated with ethanol, and stained with hematoxylin for 5 min and eosin for 3 min. Images were photographed using a 20× objective with the Eclipse C1 microscope (Nikon, Tokyo, Japan) equipped with a DS-U3 camera (Nikon, Tokyo, Japan).

### Immunofluorescence Staining

For cells, after four days of infection, the plate was fixed for 30 min with 4% paraformaldehyde (PFA). The cells were permeabilized with 0.5% TritonX-100 for 15 min. Infected cells were detected with monoclonal anti-dengue-NS3 antibody (Genetex, GTX124252) and FITC-conjugated secondary antibody (Beyotime, A0568). Finally, the cells were stained with Hoechst (Thermo Fisher Scientific, 62249) for 10 minutes and images were photographed with EVOS M5000 imaging system (Thermo Fisher Scientific, USA).

For mouse tissues, immunofluorescence staining was carried out as described previously[19]. Briefly, sections were repaired using EDTA antigen recovery buffer and incubated overnight at 4 °C with anti-dengue virus 1+2+3+4 primary antibody (Abcam, ab26837). Next, sections were incubated with CY3-conjugated secondary antibodies for 50 min and 4,6-diamidino-2-phenylindole (DAPI) for 10 min. Images were photographed using a 20× objective with the Eclipse C1 microscope (Nikon, Tokyo, Japan) equipped with a DS-U3 camera (Nikon, Tokyo, Japan).

### Transmission Electron Microscopy

The brain tissues were quickly fixed with 2.5% glutaraldehyde overnight at 4 ℃. After rinsing with PBS and fixation with 1% osmotic acid for 2 hours. Following graded acetone dehydration, tissues were infiltrated, embedded and polymerized in resin. The sections (70 nm) were stained with 2% uranyl acetate-saturated aqueous solution and lead citrate for 15 min. Imaged using a TECNAI G2 20 TWIN TEM transmission electron microscope (FEI, USA).

### Hematology

Blood (100 µl) was collected in 1 ml EDTA-K2 tubes to prevent clotting and was then briefly vortexed, and whole blood analysis was carried out using an automatic animal blood cell analyzer (TEK-VET3, China). The hematology parameters analysis included the lymphocytes, neutral counts, red and white blood cell counts, hematocrit, and platelet count.

### Cytokine detection

Expression levels of cytokines (IL-1β, IL-6, IFN-γ, TNF-α) were assayed in mouse brain tissue grindings according to the manufacturer’s instructions using different assay kits (Cloud-Clone Corp, China).

### RNA library preparation and sequencing

Total RNA was extracted from mouse brain tissue using total RNA Extractor (Trizol) according to the manufacturer’s protocol (Sangon Biotech, Shanghai). Total amounts and integrity of RNA were assessed using the RNA Nano 6000 Assay Kit of the Bioanalyzer 2100 system (Agilent Technologies, CA, USA). Then, libraries were constructed using a TruSeq Stranded mRNA LT Sample Prep Kit (Illumina, San Diego, CA, USA) according to the manufacturer’s instructions.

Libraries were sequenced on the Illumina NovaSeq 6000 platform, and 150 bp paired-end reads were generated. Index of the reference genome was built using Hisat2 (v2.0.5)[20]. The fragments per kb of transcript per million mapped reads (FPKM) was calculated using Feature Counts (v1.5.0-p3)[21].

### Identification of differentially expressed genes and functional enrichment

Using the DESeq2 R package (1.20.0), differential expression analysis of two groups (three biological replicates) was carried out. The P values were adjusted using the Benjamini & Hochberg method. For considerably differential expression, p-adj<=0.05 and |log2(foldchange)| >= 1 were chosen as the cutoff values. GO and KEGG enrichment analysis of differentially expressed genes by Clusterprofiler (3.8.1) software.

### Ethics statements and facility

All procedures involving mice and experimental protocols of our study were approved by the Animal Ethics Committee of Changchun Veterinary Research Institute. All animals were handled by the Animal Ethics Procedures and Guidelines of the People’s Republic of China, the principles described by the Animal Welfare Act, and the National Institutes of Health Guidelines for the care and use of laboratory animals in biomedical research. All animal experiments involving the dengue virus were performed in the Animal Biological Safety Level-2 laboratory (ABSL-2) of Changchun Veterinary Research Institute.

### Statistical analysis

Data are expressed as the mean ± SD values. Comparison between experimental and control groups was performed using an ordinary one-way ANOVA. Differences with a probability value of p<0.05 were defined as significant. GraphPad Prism (https://www.graphpad-prism.cn/) was used for statistical analyses and visualization. Screening for dengue virus-related genes using DisGeNET (https://www.disgenet.org)and Genecard (https://www.genecards.org) databases.

## Results

### AGB6 mice can be efficiently infected by DENV-I via intraperitoneal inoculation

To test the virulence of the DENV-I, AGB6 mice were intraperitoneally (ip.) infected with 10-fold serially diluted viral doses ranging from 10^8^ to 10^5^ TCID_50_. Observe and record the survival rate, clinical symptoms, and weight changes of mice for 15 consecutive days (Figure 1). Survival rates indicated that 10^7^ TCID_50_ and higher dose induced 100% mortality, whereas 80% and 100% survival rates were observed in animals infected with 10^6^ and 10^5^ TCID_50_, respectively (Figure 1A). By measuring the weight of mice, we found that mice infected with 10^8^ TCID_50_ experienced a rapid decrease in weight on the 2 days post-infection (dpi), ultimately leading to mouse death at 6dpi (Figure 1B); The mice infected with 10^7^ TCID_50_ showed fur wrinkles, hunchback, lethargy, Ataxia and paralysis of one or both hind limbs between 10 and 14 dpi (Figure. 1C). It is worth noting that on the 4 dpi, the trend of weight loss of mice in the 10^7^ TCID_50_ group was no different from that of mice in the 10^8^ TCID_50_ group. However, on the 7 dpi, the weight of mice in the 10^7^ TCID_50_ group slowly increased, and the status gradually recovered, until the rate of weight changes significantly decreased after the onset of paralysis symptoms, ultimately leading to death (Figure 1B). The median lethal dose (LD_50_) for DENV-I was 10^6.6^/0.5 mL, calculated by the Reed-Muench method. The dose of virus infection is positively correlated with the time of death, and heterogeneity increases with the decrease of infection dose.

**Figure 1.**
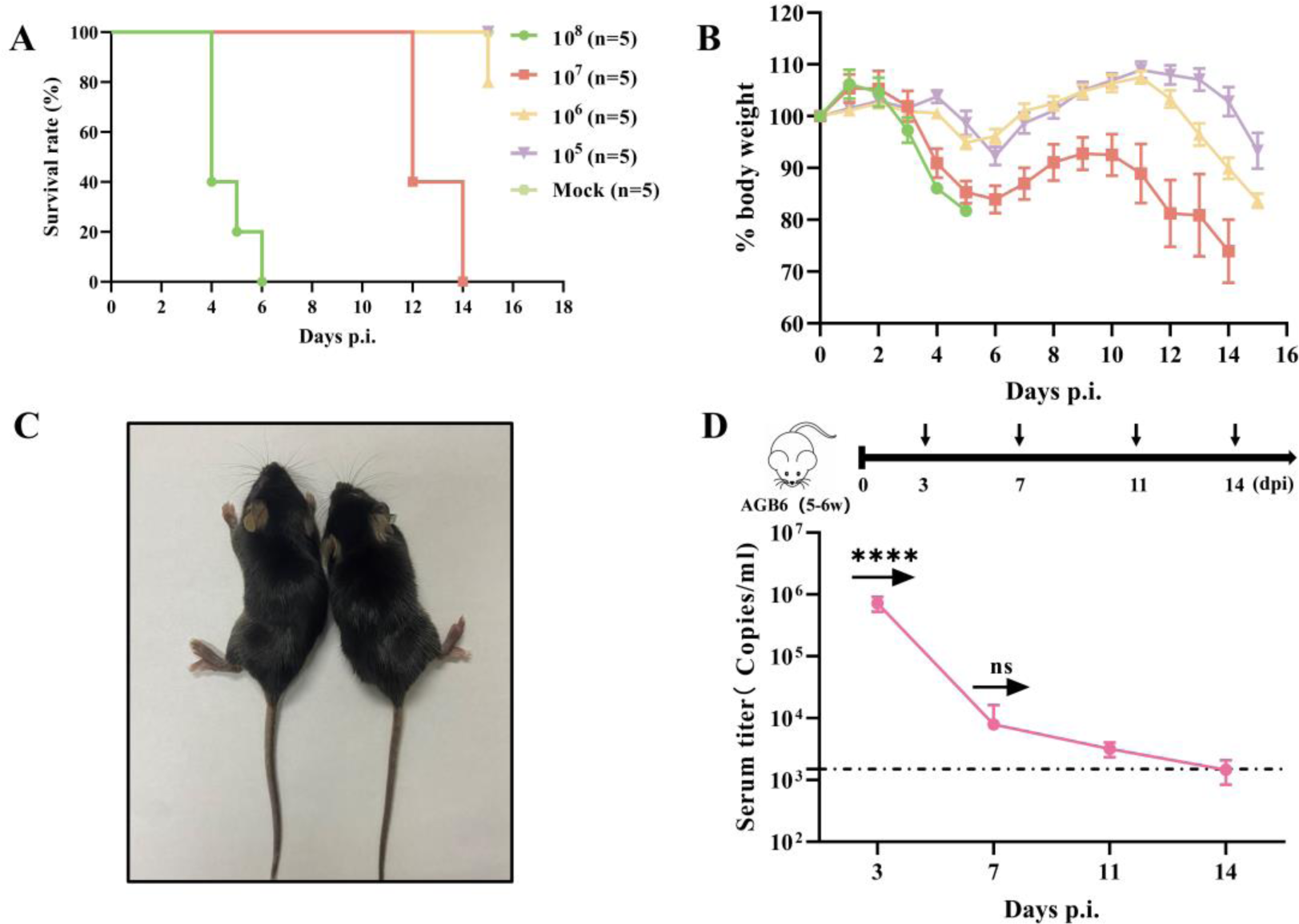
DENV-I lethal infection in AGB6 mice. **(A)** Survival probability of DENV-I challenged AGB6 mice infection via IP. routes with different titers of DENV-I. **(B)** Body weight was monitored daily. Results are expressed as the [mean ± SD] of body weight loss in percentage compared to initial body weight. **(C)** Representative images of paralysis symptoms in mice. Mice develop unilateral or bilateral hindlimb paralysis at 12dpi. **(D)** Experimental scheme. 5-6 weeks old AGB6 mice were infected with 2.5 LD_50_ of DENV-I by IP. The blood of the challenged mice was harvested (black arrow). The viremia was evaluated (n=3). The dotted black line indicates the limits of detection. Abbreviations: IP., intraperitoneally. Ordinary one-way ANOVA test. ****, p < 0.0001; ns, no significance.

Therefore, for experimental convenience and maximum stringency, a challenge dose of 2.5 LD_50_ was used in subsequent studies and monitored the virus titer in the blood during infection. The blood was collected at 3, 7, 11, and 14 dpi to detect the viral load. At 3 dpi, viremia was detected by RT-PCR, with peak loads at around 7.16×10^5^ copies/mL, followed by viral clearance from the blood circulation prior to animal death, similar to the disease kinetic described in severe dengue fever (DF) patients (Figure 1D). Furthermore, specific IgG and IgM antibodies were detected during the course of the infection, and significant IgG antibody titers were detected, which increased gradually over time, peaking at 1:51200, while the IgM antibody response peaked at 7 dpi (1:12800) and waned by 11 dpi (Supplementary Figure S1 A, B).

In addition, neither disease manifestation nor transient viremia was observed in immunocompetent BALB/c and C57BL/6 mice infected with 10^7^ TCID_50_ of DENV-I by intraperitoneal injection (IP.). (Supplementary figure S2).

### DENV-I mainly infected brain in AGB6 mice

In the previous study, we collected organs from AGB6 mice infected with DENV-I virus to evaluate viral RNA, protein expression and histopathological changes (Supplementary Figure S3). The viral protein NS1 was detected mainly in DENV-I infected mice liver at 3 dpi. After 7 dpi, the viral protein E could be detected in brain, with the highest expression during the moribund. In addition, different degrees of pathologic changes were observed in the brain, spleen, lungs and liver, and no significant pathologic changes were observed in other organs (Supplementary Figure S4).

Most importantly, it was found that DENV RNA expression was highest in brain tissue at moribund states (14 dpi). Therefore, the brain tissues were subjected to H&E staining and histopathological observation, which revealed progressive damage at both tissue and cellular levels which culminated at moribund states (Figure 2). As shown in Figure 2A, In the hippocampus of DENV-I infected mice, degeneration of neuronal cells in the granular layer (black arrows), infiltration of lymphocytes, monocytes and neutrophils (red arrows), and swelling of the vascular endothelium (yellow arrows). Cerebral cortical areas were the most severe, with neuronal cell necrosis (black arrows), perivascular inflammatory cell infiltration (red arrows), and vasodilatation (blue arrows). DENV-I infected cells were identified in neurons after staining with anti-NS3 antibody (Figure 2B). In addition, immunofluorescence staining showed clear fluorescent signals in the brains of mice at moribund states (Figure 2C). Electron microscopy analysis confirmed the presence of DENV-I particles in the brain of mice at moribund states (red arrows) (Figure 2D).

**Figure 2.**
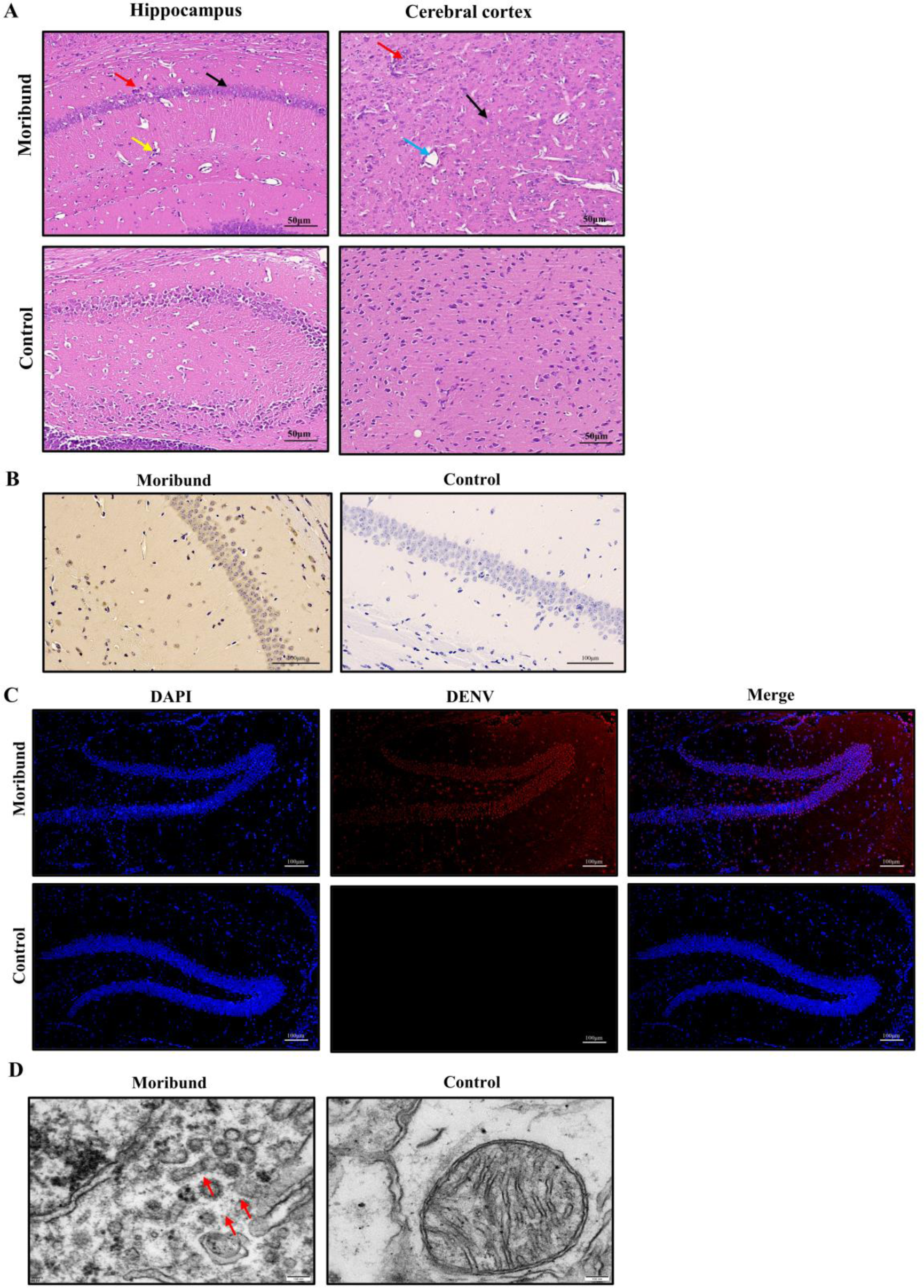
Infection status of brain tissue in DENV-I infected AGB6 mice. **(A)** Brains were harvested from IP. infected mice, and stained with H&E. Animals were euthanized at moribund states. Scale bar:50 µm. **(B)** Immunohistochemical staining of tissue sections from the DENV-I infected mice with an anti-NS3 antibody. **(C)** Immunostaining of the hippocampal at moribund states. Sections were viewed under a fluorescence microscope at 100 ×. Scale bar:100 µm. **(D)** Representative transmission electron micrographs showing viral particles in DENV-I infected mouse brain.

### Hematology in DENV-I infected AGB6 mice

Hematology has been shown to be associated with dengue-associated diseases and is tentatively used as a diagnostic and prognostic marker. To determine the effect of the dengue virus on blood chemistry in AGB6 mice, blood was collected and analyzed on 3, 7, 11, and 14 dpi. The assays included white blood cells (WBC), red blood cells (RBC), neutrophils (NEU), lymphocytes (LYM), platelets (PLT), and hematocrit (HCT) (Figure 3).

**Figure 3.**
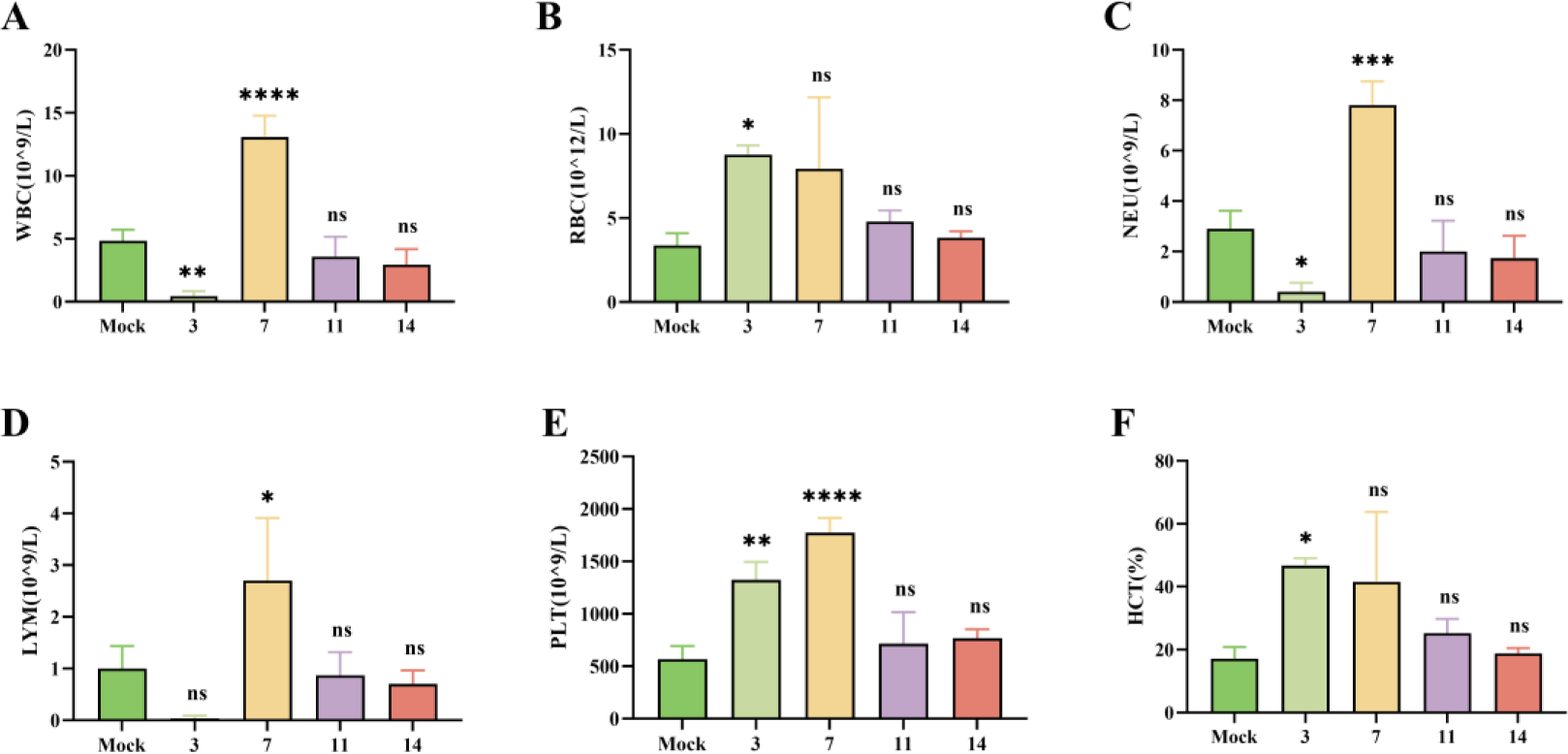
Hematology in DENV-I infected mice. AGB6 mice were IP. with 2.5 LD_50_ of DENV-I. At the indicated time points. 3 mice per group per time point were bled and euthanized. **(A)** Blood samples were processed to determine white blood cells (WBC), **(B)** red blood cells (RBC), **(C)** neutrophils (NEU), **(D)** lymphocytes (LYM), **(E)** platelets (PLT), **(F)** hematocrit (HCT) counts. Uninfected mice were used as controls. WBC, NEU, LYM, and PLT counts are given in 10^9^ cells/L, RBC counts in 10^12^ cells/L, and HCT in percentage (%). Data are expressed as the mean±SD of individual measurements. Ordinary one-way ANOVA test. *, p < 0.05; **, p < 0.01; ***, p < 0.001; ****, p < 0.0001; ns, no significance.

Compared to negative control, a significant increase in RBC, PLT concentration, and HCT was detected at 3 dpi indicating elevated blood concentrations, and conversely, at moribund states (14 dpi), the concentration decreased suggesting the occurrence of hemorrhage, notably, RBC and HCT peaked at 3 dpi (8.76 and 46.7, respectively) and PLT peaked at 7 dpi (1774) (Figure 3B, E and F). Meanwhile, a sharp decrease in the concentration of WBC, NEU, and LYM was detected during the period of high viremia (3 dpi), suggesting a possible inflammatory response of the organism, which peaked at 7 dpi as the virus was cleared from the bloodstream (13.07, 7.8 and 2.7, respectively) (Figure 3A, C and D). None of the above assays were statistically different from uninfected mice at moribund states.

In summary, hematological results indicated that DENV-I infected mice were hemorrhagic and hemoconcentrated, but thrombocytopenia (which is clinically present in 80% of patients) was not detected.

### DENV-I can be isolated from the brain tissue of infected AGB6 mice

The mice were euthanized and the brain tissue was ground in PBS buffer and the supernatant was taken for virus detection (Figure 4A). The supernatant was accessed into Vero cells, and the indirect immunofluorescence assay could observe the same fluorescence (Figure 4B) as the positive control (viral group), and the negative control had no fluorescence. Then brain tissue virus isolates were serially passaged on C6/36 cells three times. Viral RNA copy numbers and cytopathic effects were evaluated at each passage. The results showed that DENV-I isolates from the brain tissue could induce cytopathic effects (CPE) in C6/36 cells within 7 days after inoculation (Supplementary Table S2) and effectively replicated resulting in high viral RNA loads (up to 10^5^ copies/ml) (Figure 4C).

**Figure 4.**
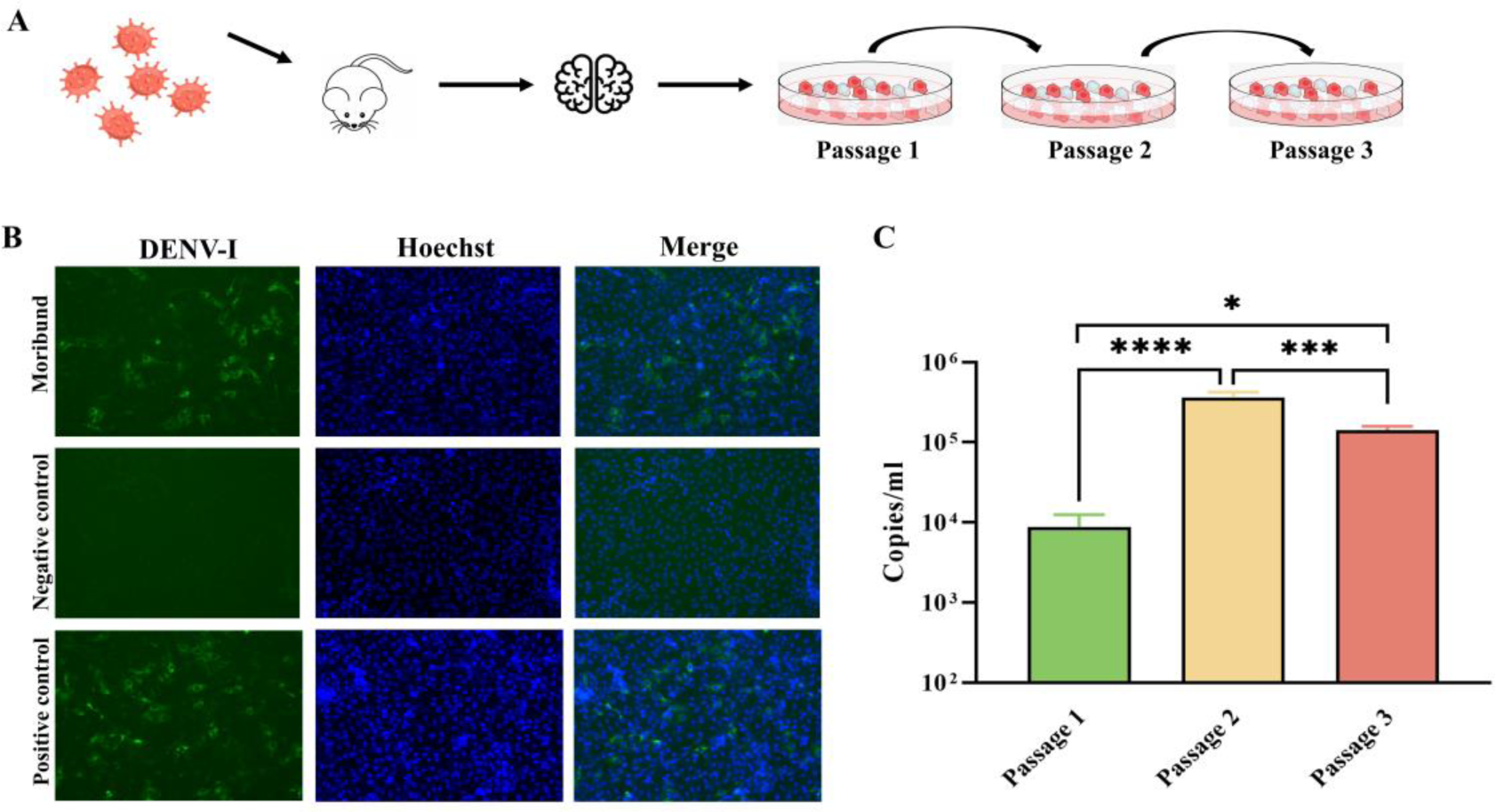
Isolation of DENV-I from C6/36 and Vero cells. **(A)** Experimental scheme. 5-6 weeks old AGB6 mice were infected with 2.5 LD_50_ of DENV-I by IP.. The brain tissues were harvested at moribund states. Isolates were passage-cultured for three generations. **(B)** Indirect immunofluorescence assay for virus detection (green fluorescent). **(C)** DENV-I was isolated from C6/36 cells and cultured for three passages; viral loads were evaluated by RT-PCR. Ordinary one-way ANOVA test. *, p < 0.05; ***, p < 0.001; ****, p < 0.0001.

### RNA sequencing and differentially expressed genes in DENV-I infected AGB6 mice brain

RNA-Seq analysis was performed using Illumina technology to compare the differences in gene expression in brain tissues of infected and healthy control mice. The samples obtained all provided complete and reasonable coverage of the different gene structures and regions of the mouse reference genome, and the high and low gene expression levels in each sample were calculated using the FPKM method (Supplementary Figure S5 and Supplementary Table S3). Both inter-sample correlation and principal component analysis (PCA) showed that there were significant differences in gene expression between the two groups, which could be used for subsequent analysis (Supplementary Fig. S6).

Subsequently, differentially expressed transcripts were screened based on the criteria of log2 (fold change) ≥ |±1| and adj. p-value < = 0.05. A total of 1,395 genes were significantly differentially expressed in the brain in dengue virus infection model mice: 1,301 upregulated and 94 downregulated genes (Supplementary figure S7A). The differentially expressed genes between the two samples were characterized by heatmaps and volcano plots (Supplementary figure S7B, C and Supplementary Table S4).

### Functional annotation of transcriptional changes

To analyze the function of differentially expressed genes (DEGs), KEGG and GO pathway enrichment analyses of the DEGs were performed using Cluster Profile software, with p-adj less than 0.05 as the threshold for significant enrichment (Figure 5A, B).

**Figure 5.**
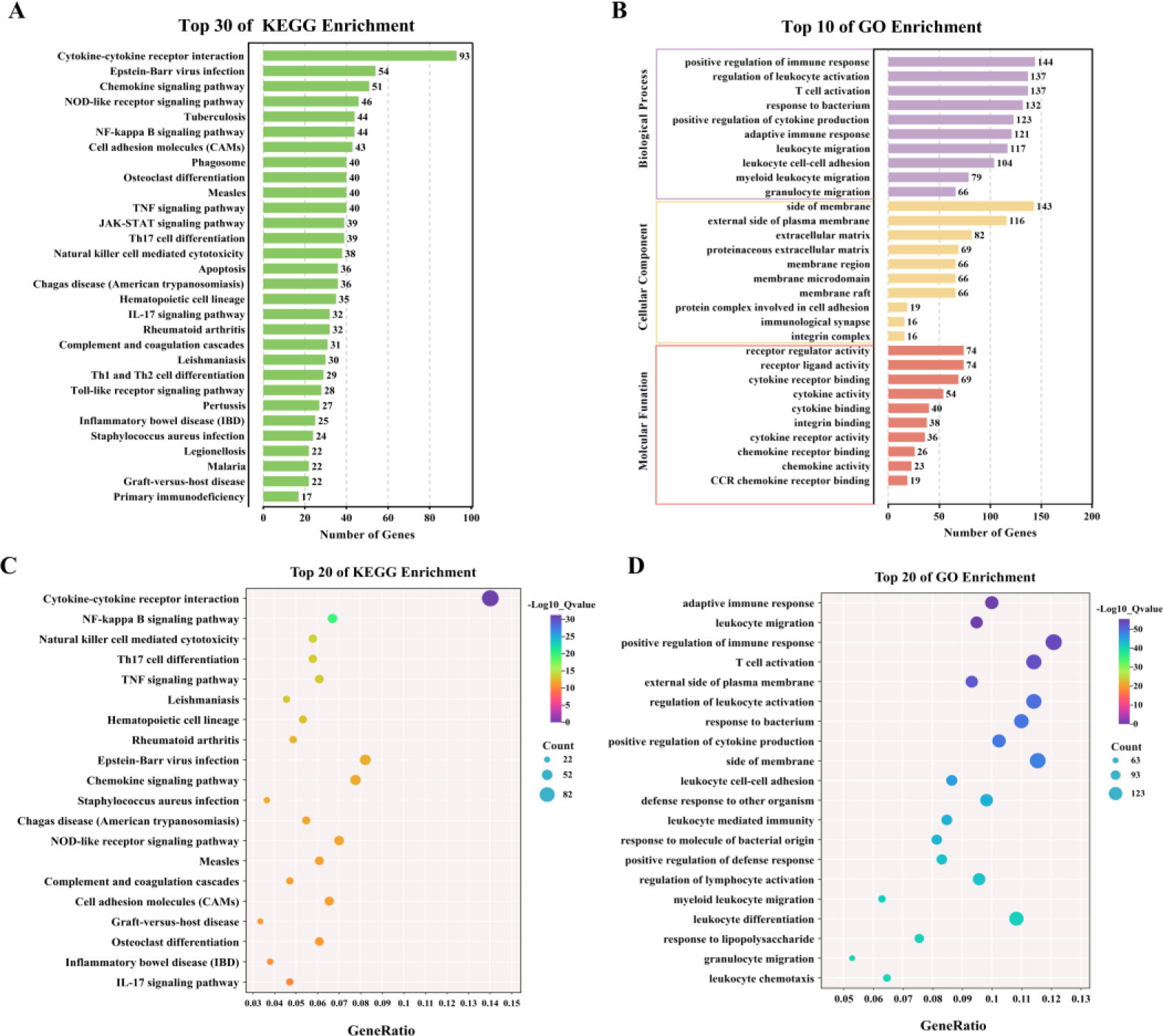
Functional Annotation of Transcriptional Changes. **(A)** KEGG enrichment analysis. **(B)** GO enrichment analysis was performed on the DEGs to describe their functions. **(C)** KEGG enrichment of gene targets of all the up-regulated mRNA. **(D)** GO enrichment of gene targets of all the up-regulated mRNA.

We then performed the KEGG pathway enrichment analysis separately for up- and down-regulated mRNA-related target genes. We found that up-regulated genes (1301 genes) were enriched for 76 significant pathways, in contrast, down-regulated genes (94 genes) were enriched none pathway (Figure 5C and Supplementary Table S5). Among them, the genes involved in Cytokine-cytokine receptor interaction, NF-kappa B signaling pathway, Natural killer cell mediated cytotoxicity, Th17 cell differentiation, the TNF signaling pathway and complement and coagulation cascades are activated after DENV infection, which involves the expression of inflammatory cytokines and the regulation of vascular permeability in the body, and these pathways have a closely related to the pathogenesis of dengue virus infection.

GO enrichment can be categorized into three parts: biological process, cellular composition, and molecular function (Figure 5D) and Supplementary Table S6). In the gene ontology, the differential genes were found to be mainly involved in the processes of immune system defense response, T cell and leukocyte activation, and cytokine-cytokine interactions when compared with the control mouse brain tissues. Taken together, the data showed that DENV infection induces a wide range of gene expression changes, some of which were involved in viral infection and others in the host’s inflammatory and immune response.

### Protein interaction network analyses and preliminary identification of brain hub genes in DENV-I infected AGB6 mice

We screened the DisGeNET and Genecard databases for genes associated with severe dengue fever and dengue shock syndrome (Supplementary Table S7 and Supplementary Table S8). After the removal of duplicated genes, 151 genes associated with severe dengue were identified by comparison with those significantly upregulated in this study. These genes were used for PPI analysis in the STRING database and Cytoscape software was used for data visualization and analysis (Figure 6A and Supplementary Table S9). We then performed tissue enrichment analysis (Supplementary Table S10). We used the MCODE plugin in Cytoscape to cluster the genes obtained from the screen (Figure 6B, C, and D) and Supplementary Table S11), which mainly included cytokines that play essential roles in the immune system, such as the Toll receptor family, chemokines, interleukins, interferons, and tumor necrosis factors. The Cytohubba insert was selected to screen for Hubba genes, the first five of which are TNF, IL-6, IL-1β, IFN-γ, and TLR4, which play essential roles in virus recognition and innate immunity.

**Figure 6.**
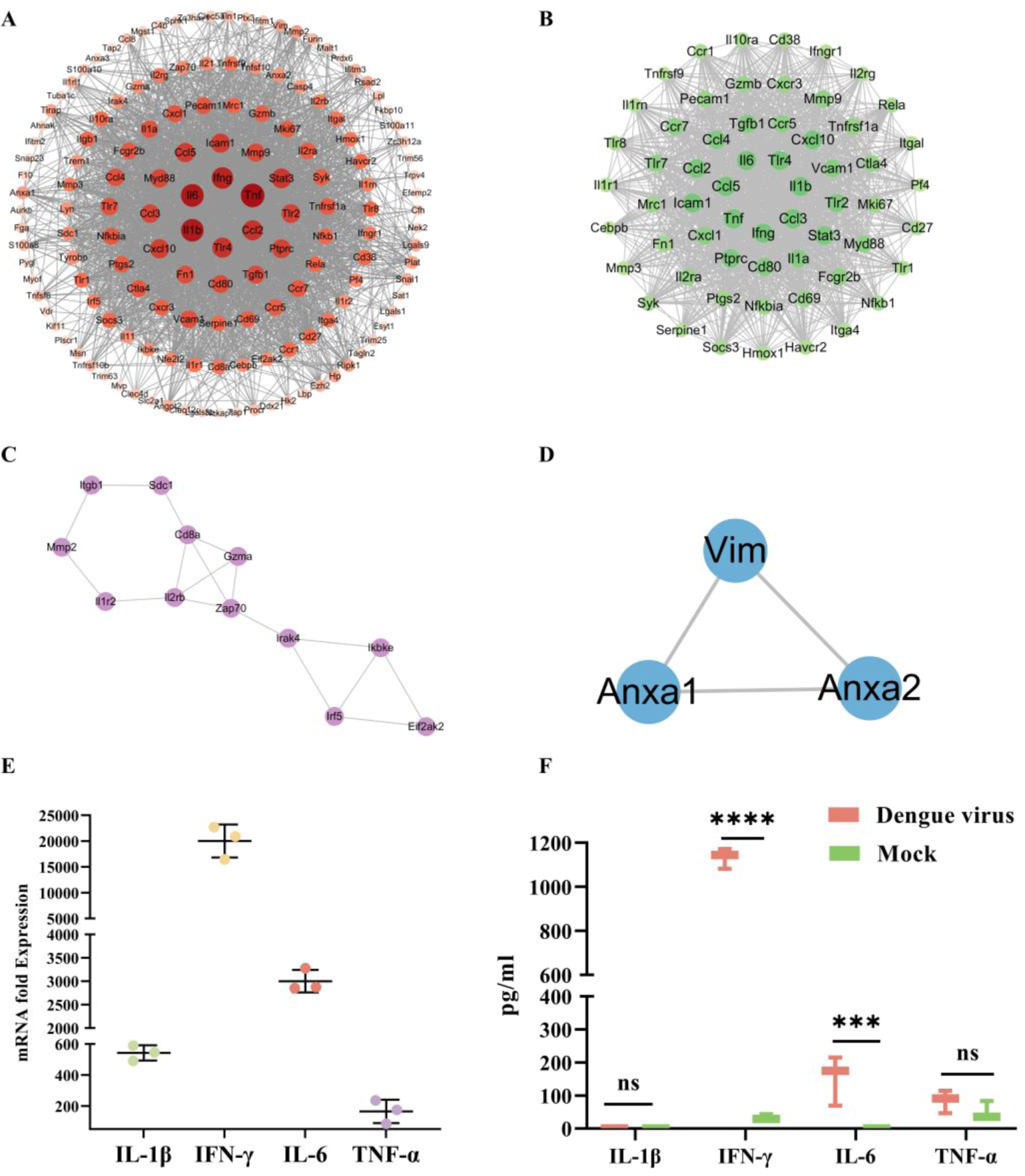
Screening for genes associated with DENV-I infection. **(A)** Network topology analysis and protein-protein interaction (PPI) networks of 151 differentially expressed genes between DENV-infected mice and controls. **(B-D)** The top three clusters that were significant from the PPI network. **(E)** Expression of pro-inflammatory cytokines in brain tissue after dengue virus infection (mRNA level). **(F)** Expression of pro-inflammatory factors in brain tissue (protein level). 2way ANOVA test. ***, p < 0.001; ****, p < 0.0001.

In order to clarify the expression of pro-inflammatory cytokines in brain tissues after viral infection, RT-PCR and ELISA were utilized for analysis, respectively. First, RT-PCR was utilized to identify these genes at the mRNA level, and the results showed that viral infection resulted in significant upregulation of these four genes, with IL-6, IL-1β and IFN-γ being the most significant (Figure 6E). Subsequently, the major chemokines (CCL2, CCL3, CCL4, and CCL5) of the CCL family members, who are also the major inflammatory chemokines in the cytokine storm, were assayed at the mRNA level (Supplementary figure S8). Finally, the changes in the protein levels of inflammatory factors in the brain tissue grinding fluid were analyzed, and the results showed that the levels of IFN-γ and IL-6 proteins in the viral group were significantly increased compared with those in the blank control group, whereas IL-1β and TNF-α did not differ from those in the control group (Figure 6F). In conclusion, we surmise that IL-6 and IFN-γ may be the key factors in dengue virus-induced neurological manifestations.

## Discussion

Dengue virus infection is one of the significant public health problems globally, especially in the tropical regions of the world, which account for 75 percent of dengue cases. Although the majority of dengue virus infections are mild or asymptomatic, about 5 percent of cases progress to severe disease, mainly attributable to sequential infections with different dengue virus serotypes[22]. For several years, the neurological manifestations of dengue virus infection have become increasingly familiar and understood and may be fatal if not treated promptly.

In 2009, the World Health Organization (WHO) issued new dengue guidelines and a new classification that includes a definition of central nervous system involvement in severe disease[23]. Currently, there are no mouse models that can be used to study dengue virus-induced central nervous system (CNS) damage. AGB6 mice are type I and type II interferon signaling pathway-deficient mice that have been used to study dengue virus transmission[1]. In this study, AGB6 mice were selected as the experimental animals, and intraperitoneal injection was used to establish a central nervous system infection model. The results of the experimental study revealed a dose-dependent dengue virus infection and infection of mice with a high dose (10^8^ TCID_50_) induced an acute lethal dengue virus infection, a viral response consistent with the study of Grace et al[18]. In contrast, infection of AGB6 mice with a low dose of the virus resulted in a transient asymptomatic systemic viral infection, followed by the death of the animal a few days after virus clearance, similar to the disease kinetics described in human[24]. Meanwhile, at 10 dpi, mice show varying degrees of observable clinical signs, including reversed coat, reduced activity, and weight loss. At 14 dpi, the moribund states, the mice deteriorate and exhibit reduced locomotion and severe manifestations of paralysis (hind limbs or unilateral hind limbs), which may lead to reduced water intake and dehydration in the animals, resulting in dramatic weight loss and ultimately death. The earliest studies in which mouse-adapted strains of DENV virus were injected into mice[25], which developed neurologic symptoms, were once thought to be unrelated to human clinical manifestations; however, the number of reports of neurologic symptoms of dengue infections is increasing[26]. AG129 mice infected intraperitoneally with DENV at a titer of 10^6^ PFU similarly exhibited neurologic abnormalities[27]. Intranasal inoculation of DENV-II at a titer of 2.4 × 10^5^ PFU in adult A6 mice resulted in systemic infection and neurologic signs[19]. The mouse-adapted strain DENV-II (D2S10) with a titer of 10^7^ PFU was infected via intraperitoneal infection of AG129 mice in a previous study without neurological symptoms (paralysis), but the presence of virus was detected in neuronal tissue[28, 29]. The strains used in the above studies were mouse-adapted strains, and it is not possible to determine whether or not the virus mutated in the mice to cause neurologic symptoms. None of the previous studies had dengue virus infection of the nervous system as a research objective, and unlike the previous studies, the strain of dengue virus used in this study was isolated from human serum (not a mouse-adapted strain), and a significant effect of high or low dosage and the duration of infection on the invasion and pathogenicity of the virus was found by observing the severity of the pathological changes in the brain tissues after the virus infection. The mouse infection model established in this study can be used to study the pathogenicity of dengue virus on the central nervous system and can be used for subsequent experimental studies.

Here, mice were infected with the virus by intraperitoneal injection, and as the number of days of infection increased, the amount of virus in the brain gradually increased, reaching a peak at moribund states. Expression of viral E protein could be detected in brain tissue after 7 dpi. Extensive pathological damage was observed in both the hippocampal region and the cerebral cortex of mouse brain tissue. Notably, the brain tissue grinding supernatant can cause the cells to show CPE phenomenon. Instead, mild damage was seen in the liver, spleen, and lungs, with NS3 antibody staining showing weak positivity. Presumably, the brain is the primary target of this strain and may follow a different mechanism of infection, further strengthening the evidence that dengue viruses infect the central nervous system. It has been shown that dengue virus causes widespread neurological involvement (e.g., transient muscle dysfunction, encephalitis), with multiple disease manifestations reflecting one or more of the following causative factors: metabolic disorders, autoimmune or excessive immune response, and direct viral invasion[30]. Encephalitis is an inflammation of the brain parenchyma, usually caused by viral infection. Direct viral invasion of the central nervous system, possibly as a result of blood-brain barrier disruption, is the proposed pathogenesis of neurologic involvement. In addition, autoimmune reactions and metabolic disorders have been shown to further worsen neurologic deterioration[31, 32]. In our study, we found that the brain tissue showed extensive pathological damage, with a large number of inflammatory cells contained within dilated cerebral blood vessels, manifesting viral encephalitis, and hematological tests revealed a mild elevation of leukocytes at 7 dpi. Furthermore, previous studies illustrate vascular leakage as a hallmark of DHF/DSS[33], leading to hemoconcentration and bleeding manifestations. Similarly, an elevated erythrocyte specific volume at 3 dpi followed by a decrease at moribund states (14 dpi) was detected in the present study. Notably, no hemorrhage was observed in pathological sections of brain tissue only vasodilatation was observed, whereas hemorrhage was observed in the liver of acutely dead mice. It was hypothesized that the strain isolated from patient sera are hypothesized to cause dengue hemorrhagic fever and encephalitis in mice, and the severity of the disease is closely related to the dose of infestation.

Hematology has been shown to be associated with dengue-associated diseases and has been used initially as a diagnostic and prognostic marker[34]. It has been shown that leukopenia often results in plasma leakage, which subsequently causes changes in capillary permeability and a massive release of cytokines or chemical mediators[35]. As large numbers of T cells are activated, a storm of cytokines is produced and acts on the vascular endothelium, which in turn manifests itself in hemorrhage, shock, and encephalopathy[36–38]. The possible mechanisms by which the virus causes the nervous system are passive transfer of CNS hemorrhage, plasma leakage or blood-brain barrier damage and infected monocyte/macrophage infiltration[30]. In this study, we found a significant decrease in leukocytes and a significant increase in erythrocyte and platelet concentrations in mice at the early stage of infection, and hematological indices suggesting an inflammatory response accompanied by hemorrhage. Infiltration of inflammatory cells is also seen in histopathologic sections of the brain. Previous studies have ruled out the possibility that lack of IFN-γ signaling due to mice impairs the thrombocytopenia mechanism[39], which required further study.

Based on transcriptomic analysis, among the significantly up-regulated genes, the major host pathways disrupted by dengue virus infection include cytokine-cytokine receptor interactions, the NF-kappa B signaling pathway, the TNF signaling pathway and complement and coagulation cascades which are thought to be involved in the pathogenesis of dengue fever closely related to the pathogenesis of dengue virus, and it is hypothesized that these factors are involved in brain tissue damage in mice. Recent studies have shown that cytokine overproduction in dengue virus infection leads to immune-mediated endothelial damage, which results in many CNS manifestations[13]. Cytokines such as tumor necrosis factor-α (TNF-α), which is usually elevated during the critical stages of dengue, may be essential cytokines in causing the disease[13]. DENV infection induces innate immunity in the host and promotes the production of pro-inflammatory factors and chemokines by immune cells, which further disrupts the blood-brain barrier (BBB) and subsequently facilitates the entry of other immune mediators into the brain, leading to neuroinflammation[40–42]. Similarly, it has been shown that NS1 antigen circulates in the bloodstream as a secreted glycoprotein that promotes cytokine release[43]. Natural killer cells are involved in the pathogenesis of neurological manifestations and activate T-helper (Th) cells, which differentiate into Th17 and Th9 cells and promote the release of pro-inflammatory cytokines into the brain to cause neuroinflammation (e.g., interferon-gamma, interleukin IL-12, IL-4, and TGF-β)[44].

In our study, brain damage in DENV I-infected mice identified by RNA-Seq was strongly associated with the release of inflammatory cytokines. IFN-β is a type I IFN and an immunoregulatory cytokine that participates in innate antiviral immune defense and regulates neuroinflammation[45, 46]. Recent studies have shown that blockage of the IFN I pathway leads to mitochondrial dysfunction in the brain, resulting in neuronal death[47]. In addition, platelets may further contribute to endothelial dysfunction through the production of interleukin 1β (IL-1β) and other inflammatory cytokines by monocytes, thereby causing neuroinflammation. It has been shown that DENV is highly sensitive to the antiviral effects of type I and II interferons (IFN)[18], and that type I and II IFNs control replication and dissemination of many viruses, including other flaviviruses[48–50]. Pretreatment of human cells with alpha/bate IFN (IFN-α/β) and IFN-γ)in a previous study protected them from DENV infection[51]. IFN-α/β receptor-mediated effects limit initial DENV replication at peripheral sites and control viral spread to the CNS, whereas IFN-γ receptor-mediated responses appear to play a role in the later stages of DEN disease by limiting viral replication in the periphery and removing virus from the CNS[52]. The AGB6 mice used in this study, which are type I and type II IFN receptor-deficient mice, were used to model the infection, and the viral load in the brain increased with the duration of infection after infection, and these results are in agreement with the published report in which the DENV-II strain (PL046), which has a titer of 10^8^ PFU, was injected into AG129 mice, and paralytic symptoms were observed, and the virus presence was detected in the nervous system[52]. Furthermore, from the data we obtained, it was found that the massive release of cytokines plays an important role in the infection of the CNS by DENV. The study found that the induction of cyclooxygenase-2 (PTGS2/COX2) by this cytokine in the central nervous system (CNS) is found to contribute to inflammatory pain hypersensitivity[47]. Equally important, IL-6, VCAM1, and ICAM1 are markers of endothelial activation, in which IL-6 triggers local inflammation by promoting leukocyte recruitment, and elevated VCAM1 and ICAM1 similarly increase endothelial cell permeability and promote the entry of cytokines in large quantities to elicit an inflammatory response[53–55].

Among the hub genes, TNF, IL-6, IL-1β, IFN-γ, and TLR4 are primarily associated with cytokine-cytokine receptor interactions. DENV infection prompts endothelial cells to release IL-6, CXCL10, and CXCL11, which increase the inflammatory response and vascular permeability, leading to plasma leakage in vivo[36]. The cytokines produced by T-cells, in addition to their role in viral clearance, promote inflammation and increase vascular permeability, which can lead to more severe disease. T-cell activation and cytokine production are observed during DHF in primary and secondary dengue virus infections [56, 57]. In severe dengue cases, a cascade of immune responses mediated by immune effectors increases the severity of the disease. Similar immune responses have been found in studies of other flaviviral infections of the nervous system, and blood-brain barrier damage induced by JEV infection was associated with increased secretion of inflammatory cytokines such as IL-1β, IL-6, and TNF-α[58]. ZIKV caused microcephaly and brain damage, and high levels of IL-1β and TNF-α were detected in the brain[59]. In order to clarify whether dengue virus-induced neurological infections follow the same mechanism of infection. In this study, high expression of IL-6, IL-1β, and IFN-γ at the mRNA level was detected by RT-PCR in brain tissues. Subsequently, the expression of inflammatory factor proteins in brain tissue was analyzed, and IFN-γ and IL-6 protein levels were significantly elevated compared to controls. It has been shown that IFN-γ and IL-6 directly affect the permeability of endothelial cells and increase the inflammatory response of the organism by inducing leukocyte recruitment, local inflammation, and damage to endothelial cells[54]. Further evidence that abnormal cytokine expression causes damage to the central nervous system.

In conclusion, the present study demonstrated that DENV can cause acute or chronic infection in AGB6 mice by intraperitoneal injection, which leads to neurological symptoms, high viral loads and significant pathological damage observed in brain tissues with clinical manifestations of paralysis. In addition, the potential mechanisms of CNS infection were initially explored by transcriptome sequencing. DENV infection in mice resulted in significant differences in the expression of a large number of genes in multiple signaling pathways in the host. Most of these differential genes (TNF, IL-6, IL-1β, IFN-γ, RELA, TLR4, VCAM1, and ICAM1) were associated with host viral defense, further demonstrating the existence of antiviral immunity in the CNS after DENV infection.

## Supportinginformation

**S1. Antibody titers in DENV-I-infected mice. (A)** Specific anti-IgG titers were determined for each serum (1:51200). **(B)** Specific anti-IgM titers were determined for each serum (1:12800).

**S2. DENV-I infection via the intraperitoneal route in C57BL/6 and BALB/c mice. (A)** Body weight was monitored daily. Results are expressed as the [mean ± SD] of body weight loss in percentage compared to initial body weight. The mice showed no disease manifestations and continued to gain weight after infection. **(B)** Blood samples were harvested and viremia was evaluated (each time point, n=3). Low viral load and mild viremia were detected in the blood. **(C)** The organs of the challenged C57BL/6 mice were harvested, and the virus titer was evaluated using RT-PCR (each time point, n=3). **(D)** The organs of the challenged BALB/c were harvested, and the virus titer was evaluated using RT-PCR (each time point, n=3). Viral loads were low in each tissue organ. The dotted black line indicates the limits of detection.

**S3. Tissue tropism and kinetic of virus replication in DENV-I infected mice. (A)** The organs (heart, liver, spleen, lung, kidney, brain, jejunum, ileum, cecum, colon, rectum) of the challenged mice were harvested, and the viral load was evaluated. **(B)** DENV viral protein expression in mouse tissue was collected at 3, 7, 11, and 14 dpi via immunoblotting with an anti-E/NS1 antibody. **(C)** Representative H&E-stained tissue sections from the spleen, liver, and lung of AGB6 mice IP. infected with 2.52 LD_50_ of DENV-I. Animals were euthanized in moribund states. Sections were viewed under a light microscope at 200 × magnifications. **(D)** Immunohistochemical staining of tissue sections from the DENV-I infected mice with an anti-NS3 antibody.

**S4. Histopathological study on Intestine, kidney, and heart. No noticeable histopathological changes were observed in these organs.**

**S5.** (A) The number of raw reads and clean reads. (B) Quality control of clean data by FastQC_30_. **(C)** The mapping rate and distribution of mapping reads to the reference genome.

**S6. Gene expression statistics of RNA-Seq**. **(A)** Boxplot of FPKM values for each independent sample. The box-and-whisker plot shows the dispersion degree of the data distribution. The horizontal axis shows the sample name, and the vertical axis shows the log_10_(FPKM+1) values. **(B)** Plot of correlation coefficients between sequencing samples. The horizontal axis shows the sample name, the right vertical axis shows the corresponding sample name, and the color represents the magnitude of the correlation coefficient. **(C)** Principal component analysis (PCA) was performed using the quantitative results of gene expression to investigate the distribution of samples.

**S7. Identification of differentially expressed genes. (A)** Statistical bar graph of differentially expressed genes. **(B)** Heatmap of differentially expressed genes. Unsupervised hierarchical clustering of differentially expressed genes. Samples from the two independent groups were assigned to the same cluster through cluster analysis. **(C)** Volcano plot of all genes in the samples from the two groups. The red dots correspond to upregulated genes, the blue dots correspond to downregulated genes, and the grey dots correspond to genes without statistically significant differences in expression.

**S8. Expression of pro-inflammatory chemokines in brain tissue after dengue virus infection (mRNA level).**

## Acknowledgments

This study was supported by Molecular Biology and Virology, Changchun Institute of Veterinary Medicine.

